# Timing of mating, reproductive status and resource availability in relation to migration in the painted lady butterfly, *Vanessa cardui*

**DOI:** 10.1101/2020.07.20.212266

**Authors:** Constantí Stefanescu, Andreu Ubach, Christer Wiklund

**Author notes:** Correspondence: C. Stefanescu, Granollers Natural Sciences Museum, Francesc Macià 51, 08402 Granollers, Spain.

## Abstract

In many migratory insects, migration occurs during the pre-reproductive phase of the life cycle. This trait probably arises from a trade-off between migration and reproduction and in females has been termed as the ‘oogenesis-flight syndrome’. However, the generality of this syndrome has been questioned, especially for monomorphic insects. We studied the relationship between migration and reproduction in the highly cosmopolitan painted lady butterfly, which in the Palaearctic undertakes the longest known multi-generational migration circuit of any insect. We tested for the oogenesis-flight syndrome in both spring and autumn migrants in two regions linked by migration, North Africa and northern Spain. Field observations were combined with laboratory experiments to determine the lifespan and the age at first mating to unravel the reproductive strategy observed in individuals captured in the wild. Females and males wait on average around 5–6 days before mating, and field data revealed that mating frequencies increase rapidly once females reach a medium wing wear category. There were seasonal differences in mating frequencies in the study regions depending on whether the region acted as a source or as a destination for migrants, and in the latter case there were almost twice as many mated females. Moreover, about 80% of females collected during migratory flights were unmated, the remaining females having mated only very recently. Our results thus strongly indicate that the painted lady fulfils the oogenesis-flight syndrome, as migration is concentrated in its relatively short pre-reproductive period. Field data also showed a high positive correlation between mating frequency and host plant abundance, which suggests that mated females have the ability to locate potential breeding areas. This, together with the very high fecundity estimated from over 1000 eggs in laboratory trials, makes the painted lady one of the most successful migratory insects on Earth.

## Introduction

Ever since the seminal works by Williams (1958) and Johnson (1969) first established that insect migration is very widespread and involves many different taxa, much research has focused on the behavioural aspects of this phenomenon. The framework developed by Kennedy (1985) defining migration as persistent or straightened individual movement, actively undertaken either through locomotion or through the use of a medium such as water or air current for transport, and temporarily suppressing otherwise routine ‘station keeping’ behaviours such as foraging, lies at the heart of this research. By adopting an individual-based approach rather than focusing on the consequences of movements at population level, entomologists tend to view migration as a behavioural phenomenon with ecological consequences (Chapman & Drake 2019). Furthermore, migration is recognised as a complex adaptation arising as a result of interactions between individuals, their genes and the environment (Cresswell et al., 2011). It is also accepted that migration is an individual attribute and is not in itself a specific trait. Within a population, a variable fraction of individuals may exhibit a *migration syndrome*, that is, a migratory phenotype combining behavioural, physiological and morphological traits that give rise to migratory movements (Dingle, 2014). For instance, in a wide range of animals including insects the migration syndrome typically includes traits such as increased food consumption coupled with fuel deposition (mainly fat) during the preparatory stage prior to migratory movements (Sapir et al., 2011).

In addition to this, one of the central tenets of the migration syndrome in insects is the so-called ‘oogenesis-flight syndrome’, which Johnson (1969) defined as the general pattern by which migratory activity in insect migrants occurs during the pre-reproductive period of the life cycle. The implicit assumption in this concept is that migration and reproduction alternate as physiological states, an idea that is reinforced by the association of migration with environmental conditions that induce adult reproductive diapause in non-migratory insects (e.g. short photoperiod or low temperatures). Even though the oogenesis-flight syndrome has been criticized by various authors on theoretical grounds and due to empirical evidence showing that some insects migrate when reproductively active (e.g. Rankin et al., 1986; Sappington & Showers, 1992; Jiang et al., 2010; Tigreros & Davidowitz, 2019), the concept is still widely accepted as a unifying principle in insect migration theory (Dingle, 2014; Chapman & Drake, 2019).

Although butterflies have played a leading role since the onset of studies of insect migration (Tutt, 1902; Williams, 1930), surprisingly little is known about the possible trade-off between migration and reproduction in this group. Work by Herman (1981), Brower (1985), Rankin et al. (1986) and Vargas et al. (2018) on the monarch butterfly, *Danaus plexippus*, has shown that the oogenesis-flight syndrome applies to migratory females during fall migration but not during spring migration since in the latter case butterflies leave their overwintering sites in Mexico having already mated and migrate with rapidly developing oocytes. Likewise, Herman & Dallmann (1981) found low levels of development in reproductive organs in painted lady butterflies, *Vanessa cardui*, collected in September-October in Minnesota, USA. Although no mention of migration was made in that paper, these butterflies were most probably migrating southwards just as they do in the equivalent season and latitudes in the western Palaearctic (Stefanescu et al., 2013). This suggests that in this species, as in the monarch butterfly, the oogenesis-flight syndrome can be applied to southward-moving migrants. However, a study by Haber (1993) calls for caution regarding the applicability of this concept in migratory butterflies since this author found that all but two of more than 250 species that were migrating eastwards across the Monteverde Cloud Forest Reserve in Costa Rica were reproductively active, carried mature eggs and spermatophores, and oviposited during migration. This study thus largely contradicts the prevailing view of a trade-off between migration and reproduction in migratory butterflies (e.g. Bhaumik & Kunte, 2018).

In this paper we examine more closely the relationship between reproduction and migration by focusing on the painted lady butterfly as a model migratory insect (Stefanescu et al., 2013). In the western Palaearctic this virtually cosmopolitan butterfly migrates from tropical Africa to Scandinavia and back again, in a succession of six or more annual generations, the longest known, regularly undertaken, multi-generational insect migration circuit (Stefanescu et al., 2013, 2016; Menchetti et al., 2019). However, although recent ecological research on the migration of this butterfly has tackled various behavioural aspects such as orientation and flight strategies (Stefanescu et al., 2007, 2013; Nesbit et al., 2009), no study has yet considered how migration and reproduction relate to each other.

Over three consecutive seasons, a large number of females were collected in two regions linked by migration, North Africa and northern Spain, to test the oogenesis-flight syndrome for both northward (i.e. spring) and southward (i.e. autumn) migration. Field observations were combined with laboratory experiments to determine the lifespan and the age at first mating in order to unravel the reproductive strategy observed in individuals captured in the wild. As pointed out by Simberloff (1981) and other authors, a good colonist needs not only to travel efficiently but also requires high fecundity and rapid reproductive development in the new habitat. We therefore also estimated the overall lifetime fecundity of females in our laboratory experiments. The painted lady is a paradigmatic example of a species establishing temporary populations in a wide range of habitats and so we thus predicted higher lifetime fecundity for females than for other related non-migratory species. Further insights into the extremely successful migratory strategy of the painted lady were obtained from an analysis of female mating frequencies in relation to the availability of egg-laying resources in the breeding region in North Africa.

## Material and methods

### 1) Age at first mating, lifetime fecundity and life span

Laboratory experiments to estimate age at first mating, lifetime fecundity and total lifespan were based on the offspring of 17 adult painted ladies, 11 males and six females, collected as wild larvae on *Cirsium arvense* growing in a gravel pit in the vicinity of Riala, Sweden (coordinates 59°37’46’’N, 18°31’23’’E). The larvae from the parental generation were reared individually in 0.5-litre plastic jars with a gauze top, on cuttings of the host plant *C. arvense*, in a room maintained at 23°C with a 22-h-light:2-h-dark cycle. In total four generations of offspring were reared.

On the day of emergence the adults were sexed, marked individually with a felt pen and transferred to a mating cage measuring 0.5×0.5×0.5 m placed under a 50-W Solar Raptor UV HID lamp. The cage contained a *Kalanchoe blossfeldiana* inflorescence on whose flowers 20 % sugar/water solution was dropped three times a day, as well as a *C. arvense* plant in a water bottle. The room in which the cages were housed was maintained on a 18-h-light:6-h-dark cycle at a temperature of 27°C during the photophase and 20°C during the scotophase. In light of previous experiences, we expected mating to begin at least 10 hours after the onset of the photophase, and so lights were turned on at 23:00 and were shut off at 05:00 the following day, and the cages were visited every 30 minutes to observe mating between 09:00 and 05:00. In this way we recorded the duration of the pre-mating period of both males and females. All matings observed were later confirmed by dissection of females post-mortem to check for the presence of a spermatophore in the female bursa copulatrix, and confirm that no females that had not been observed mating contained a spermatophore in their bursa.

We placed five males and five females in each cage on the day of emergence for all of the four offspring generations, and all individuals were allowed to live out their lives in the cage in which they had originally been placed. To measure individual female fecundity 10 mating couples were transferred to a separate cage provided with a cut specimen of *C. arvense* placed in a water bottle. A new host plant was provided daily, and the number of eggs laid were counted every day. The cages were checked daily for dead butterflies to determine individual lifespans in both males and females.

### 2) Reproductive status of wild butterflies and determinants of mating frequency

#### Sampled regions

To assess their reproductive status and its relationship with migration, wild females were collected in Morocco (Maghreb, NW Africa) and in Catalonia and the Balearic Islands (NE Spain). The Maghreb is the main source of the migrant painted ladies that colonise Spain in spring, and an important destination area in autumn for breeding butterflies of European and sub-Saharan origin (Stefanescu et al., 2011, 2016, 2017; Talavera et al., 2018; Suchan et al., 2019). Painted ladies remain in this region in low numbers throughout the winter until the spring generation emerges and emigrates to Europe. As such, the species is largely absent in the summer, when temperatures are too high (Menchetti et al., 2019).

Catalonia and the Balearic Islands are fully representative of the situation in the western Mediterranean region, the main breeding area for the painted lady in spring in the western Palaearctic migration system (Stefanescu et al., 2013). The local population that hatches emigrates soon after emergence (i.e. late May-mid June) to northern latitudes, although a few butterflies are still present in the region in the summer, especially in mountain areas. This subset of butterflies probably corresponds to a small sedentary population that does not migrate, plus some late migrants originating from southern Spain and the high mountains of North Africa. From August onwards, population levels increase again as a result of southward migration by northern European butterflies that then breed in the Mediterranean region and give rise to a small generation. Upon emergence in early autumn, these butterflies emigrate to Africa.

In Morocco, butterflies were sampled in October 2017, 2018 and 2019, in April 2018 and 2019, and in February 2018. In all, 252 females were collected to assess their reproductive status (Table 1). In Catalonia and the Balearic Islands 219 females were sampled from April–October in 2018 and 2019 (Table 1). Several sites belonging to the Catalan Butterfly Monitoring Scheme (www.catalanbms.org) were sampled repeatedly during the season and butterflies were also collected at other sites visited opportunistically.

**Table 1.** Wild females of the painted lady, *Vanessa cardui*, collected in Morocco and NE Spain to assess their reproductive status and its relationship with migration.

|  | 2017 |  | 2018 |  | 2019 |  | Total mated | Total unmated |
| --- | --- | --- | --- | --- | --- | --- | --- | --- |
|  | mated | unmated | mated | unmated | mated | unmated |  |  |
| Morocco | 9 | 9 | 48 | 64 | 51 | 71 | 108 | 144 |
| North-east Spain |  |  | 46 | 84 | 44 | 45 | 90 | 129 |
| Total | 9 | 9 | 94 | 148 | 95 | 116 | 198 | 273 |

#### Reproductive status

Once a butterfly was collected, it was sexed (based on the inspection of the tarsal structure of the front pair of legs) and its wings and body were separated and kept apart. Wings were stored in glassine envelopes and were later used for assessing wing wear (i.e. a surrogate of age) on a scale from 1 to 5 (allowing for intermediate values, e.g. 1.5), as described in Stefanescu et al. (2016).

Bodies were kept in pure ethanol in Eppendorf vials. In the case of females, however, the abdomen was separated from the thorax and kept in a second Eppendorf vial in ethanol 70% for subsequent inspection of its reproductive condition. The following four measures were then obtained: a) mating status (mated/unmated) according to the presence/absence of spermatophores in the bursa copulatrix (Burns, 1968; Svärd & Wiklund, 1989); b) number of spermatophores in the bursa copulatrix as an indicator of the number of times a female had mated; c) spermatophore condition on a scale from 1 (still unopened) to 5 (completely depleted of its content and largely broken) as an estimate of the time elapsed since mating; and d) status of the ovaries from 1 (ovarioles not immediately visible and mostly undeveloped) to 8 (ovarioles with less than 10 chorionated eggs as most eggs had already been laid) as an estimate of the developmental state of the ovaries and the proportion of the abdomen that was filled with chorionated eggs (see Appendix, Table A1 for a complete definition of the categories of spermatophore condition and ovary status).

In addition to the reproductive status of females, we also assessed visually the proportion of the abdomen that was filled with fat reserves (i.e. to find out if there was an inverse relationship between the timing of body fat depletion and ovary development). Fat content was also assessed in a sample of eight females that were reared in the laboratory to determine to what degree larval-derived resources in the form of body fat were available upon emergence.

#### Relationship between reproduction, age and migratory condition

The relationship between mating and age was modelled using logistic regression. Mating status was a binary response (0 - no spermatophore present; 1 - one or more spermatophores present in the bursa copulatrix), and wing wear was used as a surrogate of age. To establish if there was any association between reproduction and migration we tested the following two predictions:

##### Prediction 1

If the painted lady satisfies the oogenesis-flight syndrome paradigm, females will remain in a non-reproductive stage *until after* having completed migration. Therefore, mating frequency in a given region will be much lower in the period when emigrating populations emerge than when the region mainly acts as a breeding destination area.

According to this hypothesis, mating frequency in Morocco will be lower in butterflies collected in April (when the region acts as the source of migrants colonizing the Mediterranean region) than in October (when the region is a breeding destination for European and sub-Saharan migrants; see above). Likewise, in NE Spain we would expect a higher mating frequency in summer (when a large fraction of the butterflies may actually correspond to a non-migratory population) than in either spring or autumn (when most of the butterflies are immigrants arriving in the region without having mated; see above).

##### Prediction 2

If the painted lady conforms to the oogenesis-flight syndrome, females collected while they were actually migrating (i.e. undergoing straight uninterrupted flight near the ground in the expected direction, that is, northward flights in the spring; e.g. Williams, 1930, 1970; Abbot, 1951) will not have mated.

This was tested by using a subset of 20 females collected on migratory flight (17 from Morocco, 3 from Catalonia). Moreover, we also tested this second hypothesis by using an extended dataset of 72 females (48 from Morocco, 24 from Catalonia) that were collected either migrating (those used in the previous analysis) or while engaged in other activities (e.g. nectaring) on the same day and at the same locality as a conspicuous migratory wave was taking place.

In all these analyses, mating frequencies were compared using contingency tables. Data from different years were pooled to increase sample size and to reveal clearer patterns.

#### Relationship between reproduction and resource availability

If females display an oogenesis-flight syndrome, once they arrive in their destination area they will soon become reproductively active, mate and start laying eggs. However, upon arrival in a vast desert region such as much of the Maghreb, butterflies may find themselves in a very hostile environment completely devoid of resources (i.e. with no nectar or host plants; CS and AU pers. obs.). Although the hill-topping behaviour characteristically shown by the painted lady serves as an effective mechanism to secure mating rendezvous, in such cases (Shields, 1967; Rutowski, 1991) mated females will still need to locate resources that are very patchily distributed, including the nectar sources required for egg production (Hainsworth et al., 1991; Jervis et al., 2005) and host plants for egg laying. Under such a scenario, we would expect that the distribution of mated females would rapidly be biased in favour of those areas where resources are concentrated. We tested this possibility by modelling with logistic regression the relationship between mating probability and resource availability in Moroccan butterflies. We used data from autumn migrants, which arrive massively in the region in October (Stefanescu et al., 2017).

Thus, the whole of Morocco was divided into cells of 1×1 degrees of latitude/longitude (ca. 11,000 km^2^) and in each autumn survey a route of 2000-3000 km was driven that included as many different cells as possible. We tried to collect a minimum of 10 butterflies (independently of sex) from each cell and survey. In the present analysis, we use data from a subset of 18 cells for which at least 7 females had been collected (average number of females per cell was 12.89 ± 4.56).

Resource availability (i.e. nectar and host plant abundances) was assessed in each cell and survey by selecting randomly one or two sites where the cover (in m^2^) of the main host plants in the region was estimated during 60-min census. Host plants included several species of thistles (but mostly *Echinops spinosissimus* in October), common mallow (*Malva sylvestris*), desert nettle (*Forsskaolea tenacissima*) and several Boraginaceae (Stefanescu et al., 2017). A single measure of host plant abundance per cell was calculated by averaging data from individual censuses (three to six in total for each cell). Data from host plant cover were log transformed before being used in the models. Nectar availability was also assessed in each cell on a scale from 0 to 5.

The proportion of mated females in the 1 × 1-degree cells was modelled with general linear models (GLMs) using a binomial distribution and logit link function, with nectar and host plant availability as predictors. We built models using data from either all the main host plants or from only *F. tenacissima*, as we observed that this was the preferred resource for egg laying in October if present. In all, five models were built and compared using Akaike’s Information Criterion (AIC). Models that differed by *<* 2 points from the lowest AIC (ΔAIC *<* 2) were considered the top-ranked models (statistically equivalent to the best model of the set). These analyses were performed using Rstudio (R Core Team, 2018).

#### Ethical Note

The painted lady is a highly cosmopolitan species and one of the commonest butterflies all over the world, so our sampling did cause a nil impact to the populations. Our field work was licensed by the Direcció General de Polítiques Ambientals i Medi Natural de la Generalitat de Catalunya.

## Results

### Mating frequency, longevity and realised fecundity in laboratory trials

The age at first mating was estimated at 5.33 ± 1.86 (average ± SD) days in females (n= 48) and 5.22 ± 2.17 days in males (n=36). At second mating it was, respectively, 7.65 ± 2.23 (n=20) and 7.89 ± 3.96 (n=19), and at third mating, 9.67 ± 2.81 (n=6) and 7.40 ± 2.55 (n=10). There were no differences between the sexes at any of these ages (*P*>0.1 in all three cases). Seven males mated four times (at age 9.43 ± 4.0) and five males mated five times (at age 9.2 ± 2.28). The earliest age for mating was three days for females, and two days for males.

The lifespan of females was estimated at 20.29 ± 9.48 days (n=73; maximum lifespan=43 days) and for males at 20.31 ± 12.98 days (n=90; maximum lifespan=69 days, for a male that mated three times).

The mean lifetime fecundity was 1,038.1 ± 246.14 eggs (n=10 females), with a recorded maximum fecundity of 1,402 eggs. The average lifespan of this subset of females was 23.6 ± 4.95 days.

### Reproductive data for wild butterflies

Of the 471 females that were collected between 2017 and 2019, 198 had mated (42.0%) and 273 (58.0%) had not (Table 1). There was no difference in the mating frequency between Morocco (42.9%) and NE Spain (41.1%) (Chi-square test: χ^2^=0.183, *P*=0.709).

Mating status showed a very clear S-shaped relationship with wing wear, increasing very rapidly from zero for wing-wear categories of 1 and 1.5, to near 1 for wing-wear categories of 4.5 and 5 (Fig. 1). This relationship was perfectly modelled by a logistic regression, which predicted a mating probability of 0.5 at wing wear of 3.24. This result indicates that the probability of mating is strongly related to female age and that there is a critical age at which females rapidly become reproductively active and mate.

**Fig. 1.**
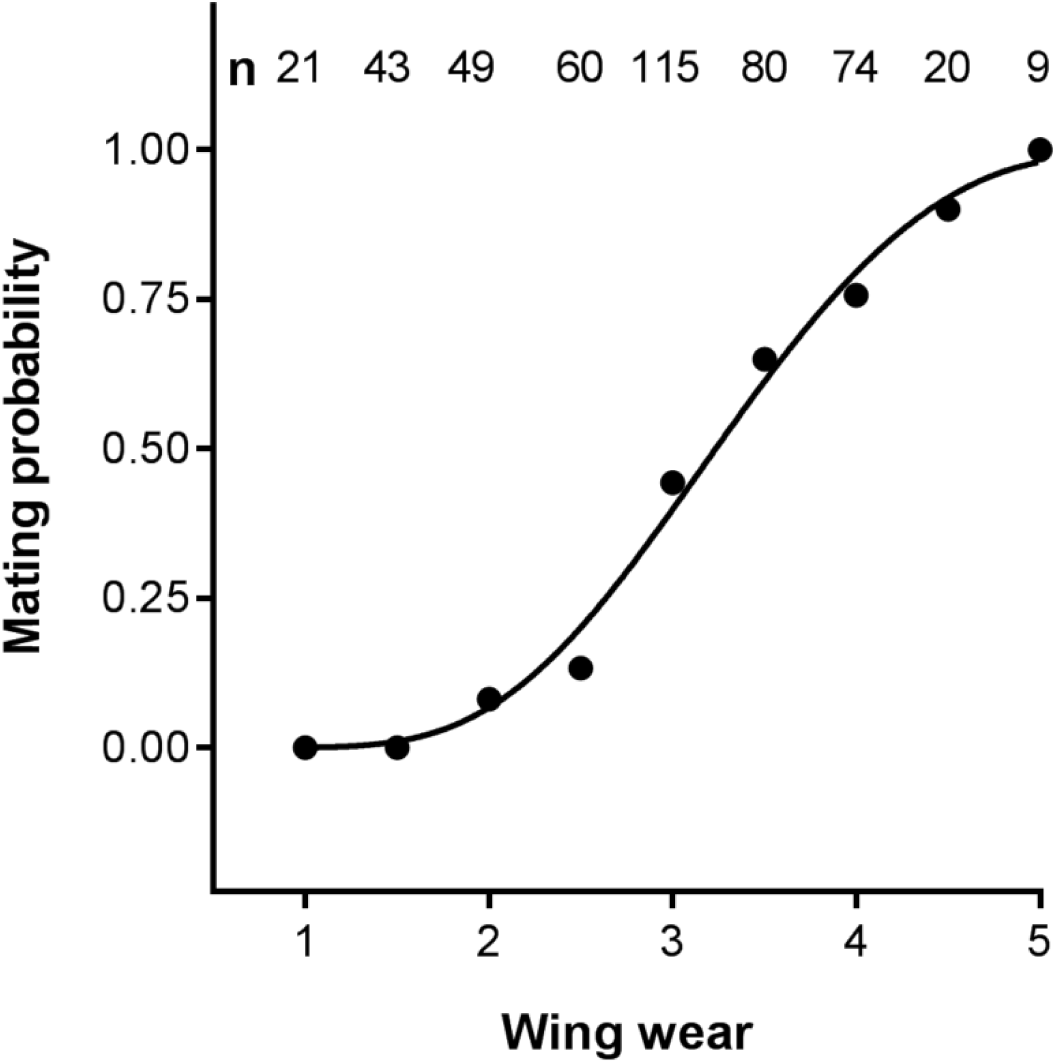
Logistic regression of mating probability against wing wear category used as a surrogate of age for 473 painted lady females collected in NE Spain and Morocco in 2017–2019. The number of females analyzed in each category is shown. Regression coefficients (± SD), z values and tests for significance in the model: Intercept: -6.4623 (± 0.6199), z value= -10.43, *P*<<0.001; wing wear: 1.9921 (± 0.1918), z value=10.39, *P*<<0.001.

Of the 198 females that had mated, 140 (70.7%) had mated once, 43 (21.7%) had mated twice, and 15 (7.6%) had mated three times. However, the average number of spermatophores per female remained similar within each category of wing wear once females had started mating (Fig. 2a). Both the status of spermatophores and ovaries followed similar S-shaped relationships with wing wear (Fig. 2b, c). On the other hand, the percentage of body fat in the abdomen showed an almost inverse relationship (Fig. 2d), with values near 100% for wing wear between 1 and 2.5, a sudden and sharp decrease for categories 3 and 3.5, and values near 0% in older butterflies (categories 4 to 5). In the eight laboratory-reared females we likewise recorded 100% body fat in the abdomen, as in freshly captured wild butterflies.

**Fig. 2.**
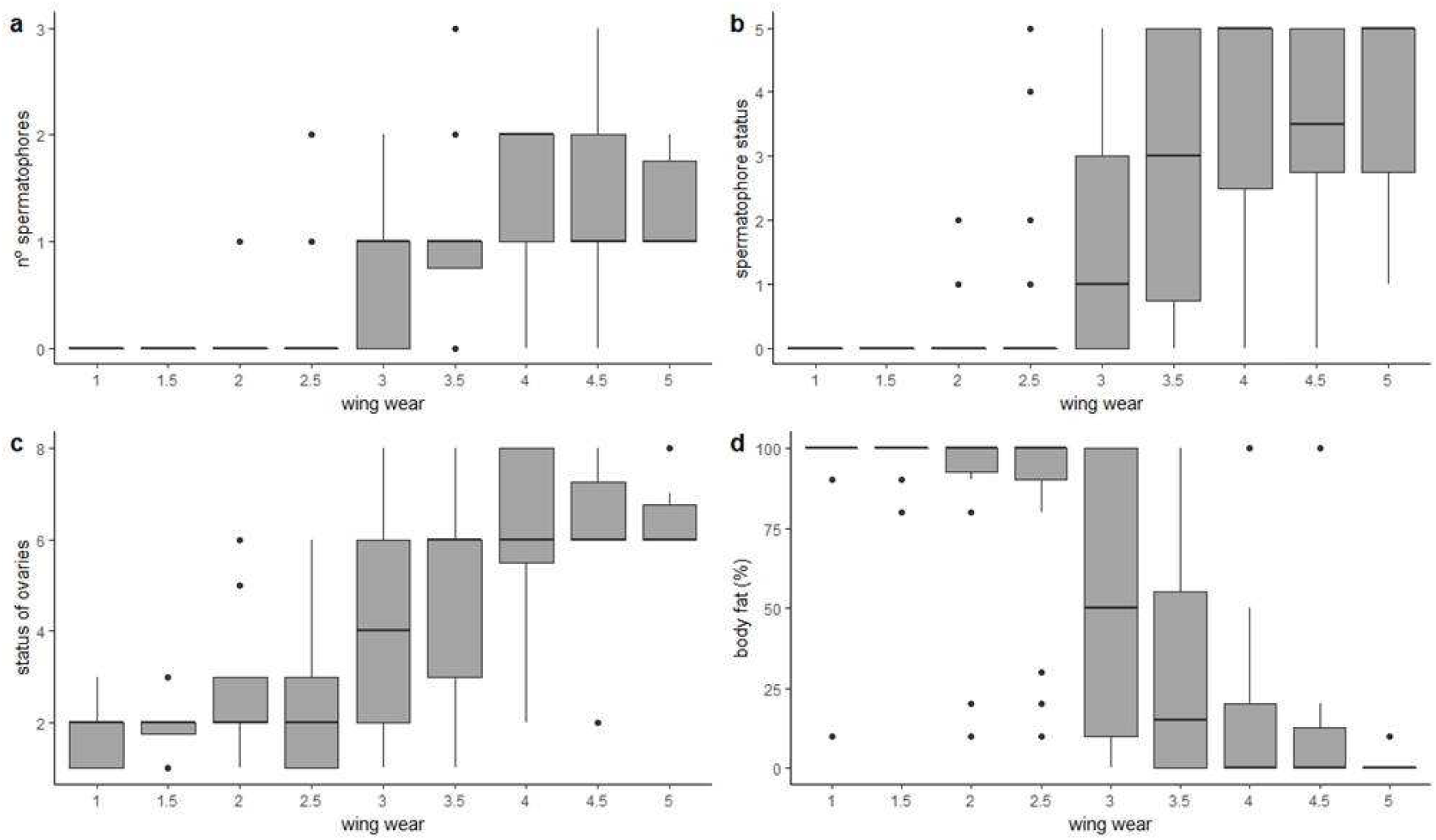
Box plots showing the relationship between wing wear and (a) number of spermatophores in the bursa copulatrix; (b) spermatophore condition; (c) status of the ovaries; and (d) proportion of body fat reserves in the abdomen. Data based on 471 painted lady females collected in NE Spain and Morocco in 2017–2019.

### Relationship between reproduction and migration

#### Seasonal differences in mating frequency in Morocco and NE Spain

A clear indication that Morocco was acting as a source of migrants in spring and a destination area for migrants in autumn is revealed by comparing the average wing wear of females (Fig. 3) and males collected in these two seasons. In April, fresh butterflies were very common in the population and overall wing wear was much lower than in October, when most butterflies in the region were immigrants. For females, the average wing wear in spring was 1.939 ± 0.998 (n=33), compared to 3.280 ± 0.715 in autumn (n=209) (F=88.903, *P*<0.0001). For males these figures were, respectively, 2.463 ± 0.911 (n=94) and 3.266 ± 0.689 (n=263) (F=78.75, *P*<0.0001).

**Fig. 3.**
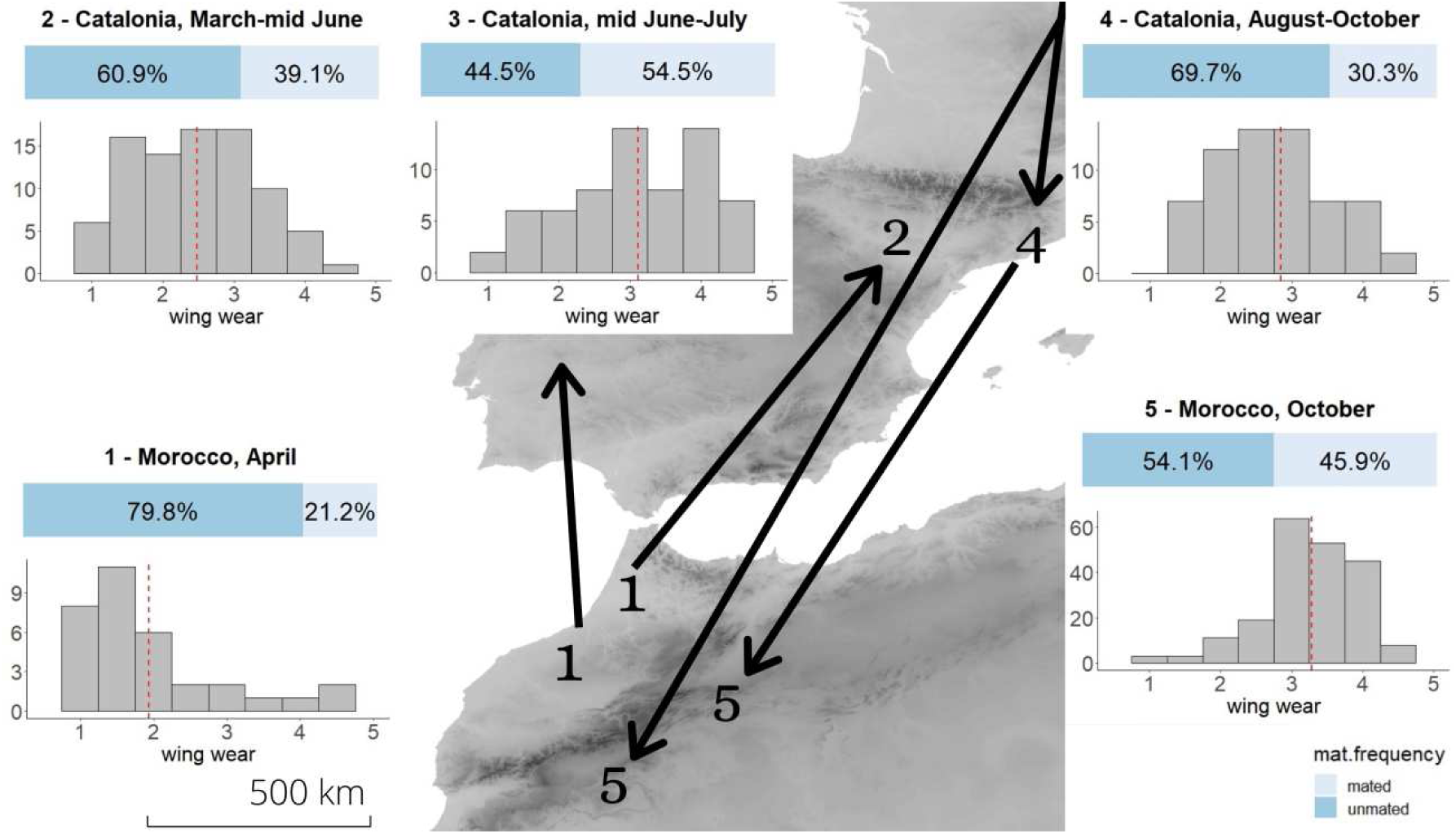
Seasonal differences in mating frequency and female population age structure (estimated from wing wear) in Morocco and NE Spain. Arrows show the main migratory seasonal movements between the study regions, with numbers at the end corresponding to the sample that was analyzed. See text for more details.

As expected under the oogenesis-flight syndrome paradigm, mating frequency in April (21.2%) was much lower than in October (45.9%) (Chi-square test: χ^2^=7.124, *P*=0.008, n=209) (Fig. 3). On the other hand, no difference in the number of spermatophores was found between both periods for *mated* females (average spermatophore counts in April and October were 1.29 and 1.30, respectively; F=0.005, *P* =0.944).

Mating frequencies in NE Spain also differed seasonally in the expected direction (χ^2^=8.254, *P*=0.016) (Fig. 3). In spring (March–mid-June) and late summer-autumn (August–October), when immigrants predominated in the population, mating frequencies were 39.1% (n=87) and 30.3% (n=66), respectively. In the summer (mid-June–July), when a large part of the population was probably composed of non-migrant butterflies, mating frequency increased to 54.5% (n=66).

#### Mating status of migrating females

Of the 20 females that were collected while on migration, 14 were unmated (70%) and six had mated (30%). All six mated females had a single spermatophore in their bursa copulatrix. The mean value for spermatophore status was 2.17, indicating that these females had mated very recently, probably the day before capture (see Appendix, Table A1).

When the sample of migratory females was extended to those butterflies that were collected at the same time and location during a notable migratory wave, 57 out of 72 females (79.2%) were unmated and 15 (20.8%) had mated. As before, all mated females had only one spermatophore in their bursa copulatrix, and the average value for spermatophore status (2.67) indicated that these females had mated recently (probably 1-2 days before capture). Mating frequency of females in this second subset was less than twice that of the remaining females (i.e. those collected on days when non-migratory flights were detected): 20.8% vs. 44% (χ^2^=13.152, *P*<0.0001).

#### Relationship between mating condition and resource availability

The best two equivalent models to account for mating frequency were those that used both host plant and nectar abundance (models 4 and 5; Table 2). Host plant abundance positively and significantly influenced mating frequency in all models where it was used as a predictor (models 1, 2, 4 and 5). The importance of the desert nettle as the primary host plant in the region in autumn is supported by equivalent models 4 and 5. On the other hand, nectar abundance did not show any significant effect on mating frequency, neither when it was used as the sole predictor (model 3) nor in combination with host plant abundance (models 4 and 5).

**Table 2.** GLM model (with binomial distribution and logit link function) testing whether or not mating frequency in Morocco in autumn is related to resource availability. Abundance of nectar sources and host plants were estimated from 3–6 random 60-min searches in 18 cells of 1×1 degrees of latitude/longitude. The two models including host plant (all species or desert nettle, *Forsskaolea tenacissima*, only) and nectar abundances were selected as the two best models.

|  | Predictors |  | Estimate | Pr(>\|z\|) | AICc | Delta |
| --- | --- | --- | --- | --- | --- | --- |
| Model 1 | Main host plants abundance (MHP) |  | 0.9431 | 6.07e-05 *** | 108.22 | -14.635 |
| Model 2 | Desert nettle abundance (DN) |  | 1.0689 | 2.29e-06 *** | 99.011 | -5.426 |
| Model 3 | Nectar abundance (N) |  | 0.1671 | 0.1665 | 108.11 | -14.525 |
| Model 4 | Main host plants + nectar abundances | MHP | 0.9024 | 0.000115 *** | 93.768 | -0.183 |
|  |  | N | 0.2418 | 0.058549 . |  |  |
| Model 5 | Desert nettle + nectar abundances | DN | 0.9049 | 0.000233 *** | 93.585 |  |
|  |  | N | 0.1635 | 0.185039 |  |  |

## Discussion

Laboratory data show that both females and males wait on average around five days before mating; field data reveal that young females with low wing wear (categories 1-2.5) have not mated and that the mating frequency increases rapidly among older females with higher wing wear (categories greater than 3.5).

The pattern of postponing mating for a number of days is unusual in butterfly groups such as Papilionidae, Pieridae and Satyrinae and few unmated females in the wild of most of the species of these groups are ever encountered (Burns, 1968; Drummond, 1984), a strategy that presumably minimizes prereproductive mortality (cf. Wiklund & Fagerström, 1977; Fagerström & Wiklund, 1982; Wiklund, 2003).

However, long waits before mating is also found in many female butterflies that hibernate as adults, which is a common trait in close nymphaline relatives of the painted lady such as the small tortoiseshell, *Aglais urticae*, the peacock butterfly, *Aglais io*, and the comma butterfly, *Polygonia c-album* (Pullin, 1986; Karlsson et al., 2008; Wiklund et al., 2019). In all of these species females typically emerge as adults but with immature ovaries without any chorionated eggs in their abdomen. This is linked to the fact that life-cycle decisions such as becoming reproductively active in the same season or delaying reproduction until the following season are made by the newly emerged adults (Wiklund et al., 2019). The painted lady and, presumably, the red admiral, *Vanessa atalanta*, neither of which have a diapause phenotype and reproduce continuously throughout the year, should nevertheless show a comparable strategy in deciding whether to migrate or not before reproduction. We found strong evidence that migration occurred before reproduction, which means that the painted lady thus conforms to the oogenesis-flight syndrome that has been proposed as the typical pattern for migratory insects. Although we do not have any results for gonad development in males to match that of ovarian development in females, the laboratory experiments on age at first mating and longevity, and also the lifetime number of matings, and, importantly, the field data on wing wear comparisons, strongly indicate that our conclusions are perfectly applicable to males. Therefore, the oogenesis-flight syndrome shown by females is well matched by a spermatogenesis-flight syndrome in males, thereby supporting our conclusion that migration usually precedes reproduction in the painted lady.

In accordance with the oogenesis-flight syndrome, the first prediction – that the mating frequency will be lower in the emergence areas that represent the source of migrants than in the destination areas where breeding occurs – was confirmed from samples in Morocco in spring and autumn. The key role of this region as the source of migrants colonizing Spain in the spring is fully supported by field work that has led to the discovery of mass emergence sites throughout the region, by multiple observations of northward migration, and by isotope and pollen markers of butterflies collected in Spain (Stefanescu et al., 2012; Talavera et al., 2018; Suchan et al., 2019). Likewise, the importance of the region as a breeding area in autumn has been confirmed by extensive field work in the last decade coupled with isotope markers (Stefanescu et al., 2016, 2017). In Morocco, not only was mating frequency in April less than half that in October but the overwhelming proportion of very fresh individuals in spring indicates that butterflies engage in migratory flights and leave the region soon after emergence before aging and wear and tear set in. The relatively short pre-reproductive period of the species (5-6 days) also means that individuals do not necessarily lose much time by migrating before the onset of reproduction. For instance, they could perfectly well migrate from Morocco to N Spain in this short pre-reproductive period if we consider that they fly downwind at ca. 45 km/h for ca. 8 h/day (e.g. by day; Stefanescu et al., 2007, 2013). Thus, a female could leave its area of emergence in Morocco in her second or third day of life, and arrive in Catalonia three days later after having travelled more than 1,000 km. Then, she would immediately mate (hill-topping activity is frenetic when migrants arrive) and start ovipositing when she is ca. one-week old.

The second prediction – that females captured while engaged in a migratory flight will not have mated – was not fully confirmed by our data. Although the majority of migratory females were unmated (about 80% for the largest dataset analysed), a small fraction had spermatophores in their bursa copulatrix. In all these cases, however, they had mated only once and the condition of the spermatophore revealed that mating had occurred very recently, at most just one or two days beforehand. This finding suggests that migratory flights are not necessarily direct but may be undertaken in different stages. The stages may serve for refuelling and building up lipid reserves used for further flight activity (as described for the monarch butterfly; Davis & Garland, 2004) but also, in the last part of the migration, for mating. Indeed, migratory females may be seen occasionally ovipositing upon arrival on host plants growing on the seashore after having crossed a large expanse of sea (CS, per. obs.).

Data from NE Spain also suggest that some painted ladies in this region are non-migrants during the summer. This can be deduced from the uninterrupted trickle of records that occur between spring and autumn migrations, and also by the higher mating frequency of summer females. Because the bulk of immigrants that arrive in the region have not mated, we would expect a larger proportion of mated females in captures from a sedentary population than in a migratory population, which indeed is what we found.

As the painted lady is one of the most cosmopolitan and successful of all insect migrants, we would also expect it to possess some biological adaptations that facilitate its ability to colonise. Laboratory trials confirmed that the most obvious of such adaptations is possibly its high fecundity compared to other butterflies. We estimated an average realised fecundity of 1,038.1 eggs, with a recorded maximum of 1,402 eggs, which almost doubles a previous estimate given by Hammad & Rafaat (1972). Our estimate places the painted lady as one of the most fecund of all butterfly species. Fecundity among Pieridae and Satyrinae varies from a low of some 100 eggs in *Leptidea sinapis* and *Lopinga achine* to figures of around 500 eggs in generalist *Pieris* species, and butterflies such as *Maniola jurtina* and *Hipparchia semele* (Wiklund & Karlsson, 1988; Wiklund et al., 2001). For the comma butterfly, a close relative of the painted lady, lifetime fecundity lies in the range 300–400 in the hibernating generation, and 500–600 in the directly developing generation (Karlsson et al., 2008), while in the migratory monarch butterfly fecundity lies in the range 600–700 (Svärd & Wiklund, 1988). On the other hand, the lifespan of the painted lady lies in a range of 3-4 weeks, typically recorded for many other butterfly species (Jervis et al., 2005). From an ecological viewpoint the exceptionally high fecundity of the painted lady means that it has a excellent potential for population growth, which helps explain the high numbers of individuals that can be seen during migration in years in which environmental conditions are favourable (e.g. Stefanescu et al., 2013; Benyamini, 2017). Interestingly, higher fecundity consistent with that found in the painted lady has also been recorded in migratory noctuid moths relative to other related non-migratory species (Spitzer et al., 1984). More generally, high fecundity is a common pattern in migratory insects and other animals that counterbalances mortality associated with migration (Sibly et al. 2012; Chapman et al., 2015) and can be regarded as a characteristic trait and integral part of the dispersal syndrome (Stevens et al., 2014).

The relatively late egg maturation that we determined for the painted lady also contributes to its successful migratory strategy. Within the framework developed by Jervis et al. (2005), the painted lady has an ovigeny index of zero (i.e. the number of eggs females have ready to lay divided by the lifetime potential fecundity), which means that resources derived from larval feeding can be used entirely in building up lipid reserves to fuel the migratory flight. This was confirmed in eight reared females that, upon emergence, had a full body fat reserve (e.g. an abdomen completely filled with fat), as has been found in the monarch butterfly (Brown & Chippendale, 1974). For the monarch butterfly, Gibo & McCurdy (1993) calculated that full body fat can power about 40 hours of flight. Although no similar study is available for the painted lady, it seems highly likely that a full body fat reserve may similarly allow for long migratory flights even in the absence of additional resources derived from adult feeding at the emergence sites.

Finally, as we expected, host plant availability is a good predictor of mating frequency and hence mated females are commoner in areas where host plants are abundant. On the other hand, nectar abundance did not explain mating frequency, even if it has been shown to be the main predictor of adult abundance in autumn in Morocco (Stefanescu et al., 2017). This suggests that when they arrive in the region, females (and males) first concentrate in areas that are rich in nectar sources. For females, nectar represents the essential energy source to be converted into egg production (Hainsworth et al., 1991), while for males it is equally important for sustaining the highly demanding hill-topping behaviour. That encounter-sites for mating in the painted lady are not necessarily associated with the presence of nectar sources but, rather, with topographic features such as hilltops (cf. Shields, 1967; Rutowski, 1991) is confirmed by the lack of predictive power of nectar availability on mating frequency (Table 2). On the other hand, the high positive effect of host plant abundance on mating frequency strongly suggests that mated females have developed a great ability to locate potential breeding areas. Indeed, we have many observations of host plants occurring in extreme isolation in the middle of vast extensions of desert that had been located by females and had received large number of eggs. This capacity, in combination with the very high fecundity already mentioned and one of the broadest diets known for any butterfly (Celorio-Mancera et al., 2016), makes the painted lady an exceptionally good colonist and one of the most successful migratory insects on Earth.

## Acknowledgements

We are very grateful to the following people who helped in the field work: Antoni Arrizabalaga, Carlos Hernández-Castellanos, Pau Colom, Oriol Massana, Juli Mauri, Marta Miralles. Local people in Morocco were always very kind and helped in many ways. Rhonda Snook and Karl Gotthard kindly arranged a stay by CS to work with CW in Stockholm University. Jason Chapman and Don Reynolds commented on a previous version. Mike Lockwood revised the English version. Our work was partially funded by Fundació Barcelona Zoo. We declare we have no competing interests.

## Appendix Table A1

Categories used for classifying spermatophore condition and ovary status. Spermatophore condition was used to assess the time elapsed since mating, while ovary status was used as an estimate of the developmental state and the proportion of the abdomen that was filled with chorionated eggs.

### (1) Spermatophore condition

1. spermatophore unopened (female had mated just before being collected, probably the same day).
2. spermatophore in the process of being opened by the lamina dentata of the inner wall of the bursa copulatrix (female had mated not long before being collected, probably the day before).
3. spermatophore had been opened by the lamina dentata but only recently (it was still rounded but its outer envelope had already two scars produced by the lamina dentata; mating had occurred about two days before the female was collected).
4. spermatophore was about half full, the outer wall having disappeared from the bursa copulatrix (female had mated about three days before being collected).
5. spermatophore was completely depleted of its content and only the outer wall remained (female had mated several days before being collected).

### (2) Status of the ovaries

1. ovarioles were so small that they were not immediately visible, which means that ovaries were in their least developed stage.
2. very small immature oocytes present in the *germarium* region of the ovarioles.
3. oocytes still growing as they were in the process of being yolked.
4. abdomen filled with non-chorionated eggs.
5. abdomen filled with eggs, 50% of which were non-chorionated and 50% chorionated.
6. abdomen was completely full of chorionated eggs.
7. the abdomen was half empty because a large proportion of eggs had already been laid.
8. there were less than 10 chorionated eggs remaining in the abdomen, indicating that the female had laid most of her eggs.

### Bullet points

1. Females and males wait on average around 5–6 days before mating
2. Migration is concentrated in the relatively short pre-reproductive period
3. The painted lady is one of the most fecund of all butterfly species
4. Mated females have an exceptional ability to locate potential breeding areas
5. These traits conform a migration syndrome and allow a successful migratory strategy

## Notes

### Competing Interest Statement

The authors have declared no competing interest.

## References

Abbot, C.H. (1951) A quantitative study of the migration of the painted lady butterfly, *Vanessa cardui* L. Ecology, 32, 155–171.

Benyamini, D. (2017). A swarm of millions of *Vanessa cardui* (Linnaeus, 1758) in winter-spring 2015-2016 in the south-east Mediterranean - The missing link (Lepidoptera, Nymphalidae). Atalanta, 48, 103–128.

Bhaumik, V., & Kunte, K. (2018). Female butterflies modulate investment in reproduction and flight response to monsoon-driven migrations. Oikos, 127, 285–296.

Brower, L.P. (1985). New perspectives on the migration biology of the monarch butterfly, *Danaus plexippus* L. In M.A. Rankin (Ed.), Migration: mechanisms and adaptive significance (pp. 748-785), Contributions in Marine Sciences Supplement, vol. 27. Port Aransas, Texas: The University of Texas.

Brown, J.J., & Chippendale, G.M. (1974). Migration of the monarch butterfly, *Danaus plexippus*: energy sources. Journal of Insect Physiology, 20, 1117–1130.

Burns, J.M. (1968). Mating frequency in natural populations of skippers and butterflies as determined by spermatophore counts. Proceedings of the Natural Academy of Sciences USA, 61, 852–859.

Celorio-Mancera, M. de la P., Wheat, C.W., Huss, M., Vezzi, F., Neethiraj, R., Reimegard, J., Nylin, S., & Jan, N. (2016). Evolutionary history of host use, rather than plant phylogeny, determines gene expression in a generalist butterfly. BMS Evolutionary Biology, 16, 59 (2016).

Chapman, J.W., & Drake, V.A. (2019). Insect migration. In M.D. Breed & J. Moore (Eds.), Encyclopedia of Animal Behavior, 2n ed. (pp. 573–580). Oxford: Academic Press.

Cresswell, K.A., Satterthwaite, W.H., & Sword, G.A. (2011). Understanding the evolution of migration through empirical examples. In E.J. Milner-Gulland, J.M. Fryxell & A.R.E. Sinclair (Eds.), Animal migration. A synthesis (pp. 7–16). Oxford: Oxford University Press.

Davis, A.K., & Garland, M.S. (2004). Stopover ecology of monarchs in coastal Virginia: using ornithological techniques to study monarch migration. In K.S. Oberhauser & M.J. Solensky (Eds.), The monarch butterfly. Biology and coservation (pp. 89–96). New York: Cornell University Press.

Dingle, H. (2014). Migration. The biology of life on the move. Oxford: Oxford University Press.

Drummond III, B.A. (1984). Multiple mating and sperm competition in the Lepidoptera. In R.L. Smith (Ed.), Sperm competition and the evolution of animal mating systems (pp. 291–370). Orlando: Academic Press.

Fagerström, T., & Wiklund, C. (1982). Why do males emerge before females? Protandry as a mating strategy in male and female butterflies. Oecologia, 52, 164–166.

Gibo, D.L., & McCurdy, J. (1993). Lipid accumulation by migrating monarch butterflies (*Danaus plexippus* L.). Canadian Journal of Zoology, 71, 76–82.

Haber, W.A. (1993). Seasonal migration of monarchs and other butterflies in Costa Rica. Natural History Museum of Los Angeles County, Science Series, 38, 201–207.

Hainsworth, F.R., Precup, E., & Hamill, T. (1991). Feeding, energy processing rates and egg production in painted lady butterflies. Journal of experimental Biology, 156, 249–265.

Hammad, S.M., & Raafat, A.M. (1972). The biology of the painted lady butterfly, *Vanessa (Pyrameis) cardui* L. (Lepidoptera: Nymphalidae). Bulletin de la Société Entomologique d’Égypte, 56, 15–20.

Herman, W.S. (1981). Studies on the adult reproductive diapause of the monarch butterfly, *Danaus plexippus*. Biological Bulletin, 160, 89–106.

Herman, W.S., & Dallmann, S.H. (1981). Endocrine biology of the painted lady butterfly *Vanessa cardui*. Journal of Insect Physiology, 27, 163–168.

Jervis, M.A., Boggs, C.L., & Ferns, P.N. (2005). Egg maturation and its associated trade-offs: a synthesis focusing on Lepidoptera. Ecological Entomology, 30, 359–375.

Jiang, X.F., Luo, L.Z., & Sappington, T.W. (2010). Relationship of flight and reproduction in beet armyworm, *Spodoptera exigua* (Lepidoptera: Noctuidae),a migrant lacking the oogenesis-flight syndrome. Journal of Insect Physiology, 56, 1631–1637.

Johnson, C.G. (1969). Migration and dispersal of insects by flight. London: Methuen.

Karlsson, B., Stjernholm, F., & Wiklund, C. (2008). Developmental trade-offs in a polyphenic butterfly: lifespan versus reproduction. Functional Ecology, 22, 121–126.

Kennedy, J.S. (1985). Migration: behavioral and ecological. In M.A. Rankin (Ed.), Migration: mechanisms and adaptive significance (pp. 5-26). Contributions in Marine Sciences Supplement, vol. 27. Port Aransas, Texas: University of Texas.

Menchetti, M., Guéguen, M., & Talavera, G. (2019). Spatio-temporal ecological niche modelling of multigenerational insect migrations. Proceedings of the Royal Society Series B. Biological Sciences, 286, 20191583.

Nesbit, R.L., Hill, J.K., Woiwod, I.P., Sivell, D., Bensusan, K.J., & Chapman, J.W. (2009). Seasonally adaptive migratory headings mediated by a sun compass in the painted lady butterfly, *Vanessa cardui*. Animal Behaviour, 78, 1119–1125.

Pullin, A.S. (1986). Effect of photoperiod and temperature on the life-cycle of different populations of the peacock butterfly *Inachis io*. Entomologia experimentalis et applicata, 41, 237–242.

R Core Team (2018) R: A language and environment for statistical computing. R foundation for statistical computing, Vienna, Austria. https://www.R-project.org/.

Rankin, M.A., McAnelly, M.L., & Bodenhamer, J.E. (1986). The oogenesis-flight syndrome revisited. In W. Danthanarayama (Ed.), Insect flight: dispersal and migration (pp. 27–88). Berlin-Heidelberg: Springer.

Rutowski, R.L. (1991). the evolution of male mate-locating behavior in butterflies. The American Naturalist, 138, 1121–1139.

Sapir, N., Butler, P.J., Hedenström, A., & Wikelski, M. (2011). Energy gain and use during animal migration. In E.J. Milner-Gulland, J.M. Fryxell & A.R.E. Sinclair (Eds.), Animal migration. A synthesis (pp. 52–67). Oxford: Oxford University Press.

Sappington, T.W. & Showers, W. (1992). Reproductive maturity, mating status, and long-duration flight behavior of *Agrotis ipsilon*, and the conceptual misuse of the oogenesis-flight syndrome by entomologists. Environmental Entomology, 21, 677–688.

Shields, O. (1967). Hilltopping: An ecological study of summit congregation behavior of butterflies on a south California hill. Journal of Research on the Lepidoptera, 6, 69–178.

Sibly, R.M., Witt, C.C., Wright, N.A., Venditti, C., Jetz, W., & Brown, J.H. (2012). Energetics, lifestyle, and reproduction in birds. Proceedings of the Natural Academy of Sciences USA, 109, 10937–10941.

Simberloff, D. (1981). What makes a good island colonist? In R.F. Denno & H. Dingle (Eds.), Insect life history patterns: habitat and geographic variation (pp. 195–205). Berlin-Heidelberg: Springer.

Spitzer, K., Rejmánek, M., & Soldán, T. (1984). The fecundity and long-term variability in abundance of noctuid moths (Lepidoptera, Noctuidae). Oecologia, 62, 91–93.

Stefanescu, C., Alarcón, M., & Ávila, A. (2007). Migration of the painted lady butterfly, *Vanessa cardui*, to north-eastern Spain is aided by African wind currents. Journal of Animal Ecology, 76, 888–898.

Stefanescu, C., Alarcón, M., Izquierdo, R., Paramo, F., & Ávila, A. (2011). Moroccan source areas of the painted lady butterfly *Vanessa cardui* (Nymphalidae: Nymphalinae) migrating into Europe in spring. Journal of the Lepidopterist Society, 65, 15–26.

Stefanescu, C., Páramo, F., Åkesson, S., Alarcón, M., Ávila, A., Brereton, T., Carnicer, J., Cassar, L.F., Fox, R., Heliöla. J., …, & Chapman, J. (2013). Multi-generational long distance migration of insects: studying the painted lady butterfly in the Western Palaearctic. Ecography, 36, 474–486.

Stefanescu, C., Puig-Montserrat, X., Samraoui, B., Izquierdo, R., Ubach, A., & Arrizabalaga, A. (2017). Back to Africa: autumn migration of the painted lady butterfly *Vanessa cardui* is timed to coincide with an increase in resource availability. Ecological Entomolology, 42, 737–747.

Stefanescu, C., Soto, D.X., Talavera, G., Vila, R., & Hobson, K.A. (2016). Long distance autumn migration across the Sahara by painted lady butterflies: exploiting resource pulses in the tropical savannah. Biology Letters, 12, 20160561.

Stevens, V.M., Whitmee, S., Le Galliard, J.-F., Clobert, J., Böhning-Gaese, K., Bonte, D., Brändle, M., Dehling, D.M., Hof, C., Trochet, A., & Baguette, M. (2014). A comparative analysis of dispersal syndromes in terrestrial and semi-terrestrial animals. Ecology Letters, 17, 1039–1052.

Suchan, T., Talavera, G., Sáez, L., Ronikier, M., & Vila, R. (2019). Pollen metabarcoding as a tool for tracking long-distance insect migration. Molecular Ecology Resources, 19, 149–162.

Svärd, L., & Wiklund, C. (1988). Fecundity, egg weight and longevity in relation to multiple mating in females of the monarch butterfly. Behavioral Ecology and Sociobiology, 23, 39–43.

Svärd, L., & Wiklund, C. (1989). Mass and production rate of ejaculates in relation to monandry/polyandry in butterflies. Behavioral Ecology and Sociobiology, 24, 395–402.

Talavera, G., Bataille, C., Benyamini, D., Gascoigne-Pees, M., & Vila, R. (2018). Round-trip across the Sahara: Afrotropical painted lady butterflies recolonize the Mediterranean in spring. Biology Letters, 14, 20180274.

Tigreros, N., & Davidowitz, G. (2019). Flight-fecundity tradeoffs in wing-monomorphic insects. Advances in Insect Physiology, 56, 1–41.

Tutt, J. W. (1902). The migration & dispersal of insects. Elliot Stock, London.

Vargas, N.R., Rowe, L., Stevens, J., Armagost, J.E., Johnson, A.C., & Malcolm, S.B. (2018). Sequential partial migration across monarch generations in Michigan. Animal Migration, 5, 104–114.

Wiklund, C. (2003). Sexual selection and the evolution of butterfly mating systems. In P.R., Ehrlich, W.B. Watt, & C. Boggs, C. (Eds.), Butterflies, Ecology and Evolution taking flight (pp. 67–90). Chicago: The Chicago University Press.

Wiklund, C., & Fagerström, T. (1977). Why do males emerge before females? A hypothesis to explain the incidence of protandry in butterflies. Oecologia, 31, 153–158.

Wiklund, C., & Karlsson, B. (1988). Sexual size dimorphism in relation to fecundity in some Swedish satyrid butterflies. American Naturalist, 131, 132–138.

Wiklund, C., Karlsson, B., & Leimar, O. (2001). Sexual conflict and cooperation in butterfly reproduction – a comparative study on polyandry and female fitness. Proceedings of the Royal Society Series B. Biological Sciences, 268, 1661–1667.

Wiklund, C., Lehman, P., & Friberg, M. (2019). Diapause decision in the small tortoiseshell butterfly, *Aglais urticae*. Entomologia Experimentalis et Applicata, 167, 433–441.

Williams, C.B. (1930). Migration of butterflies. Oliver and Boyd, Edinburgh.

Williams, C. B. (1958). Insect migration. Collins, London.

Williams, C.B. (1970). The migrations of the painted lady butterfly *Vanessa cardui* (Nymphalidae), with special reference to North America. Journal of the Lepidopterist Society, 24, 157–175.

